# Variables affecting acquisition and maintenance of operant ethanol self-administration in male and female Long-Evans rats

**DOI:** 10.1101/2024.02.29.581642

**Authors:** Shannon R Wheeler, Joseph R Pitock, Arleen Perez Ayala, Shikun Hou, Nathaly M Arce Soto, Elizabeth J Glover

## Abstract

**Aims:** The goal of the present study was to determine the effect of prior experience with ethanol drinking and changes in session duration on the acquisition and maintenance of operant ethanol self-administration.

**Methods:** Adult male and female Long-Evans rats were trained to operantly self-administer ethanol. A subset of male rats underwent intermittent-access two-bottle choice drinking in the home cage prior to operant training. Controls were given access to two bottles of water. Once fully trained in 30-min operant sessions, session duration was reduced to 15 min for all male and female rats. Differences between 30- and 15-min sessions were also assessed in a separate group of male and female rats trained to self-administer sucrose.

**Results:** No differences were observed in acquisition rates, the magnitude of responding for ethanol, or total ethanol consumed between male rats allowed to drink ethanol in the home cage and those that remained ethanol naïve prior to operant training. A significant decrease in appetitive and consummatory behaviors was observed in males trained to lever press for either ethanol or sucrose. Females exhibited a similar decrease in operant performance for sucrose, but their behavior was largely unchanged in response to changes in session duration when ethanol was the reinforcer.

**Conclusions:** These data suggest that the use of prior home cage ethanol drinking as an initiation procedure offers little advantage over no initiation procedure at all. Moreover, reducing operant session duration from 30-min to 15-min has the potential to decrease, rather than increase, levels of ethanol intake.

**Short summary:** Intermittent-access two-bottle choice ethanol drinking offers no advantage as an initiation procedure for operant ethanol self-administration over animals that are ethanol-naïve prior to training. In addition, shortening the operant session duration does not increase overall intake or promote binge-like patterns of intake for either ethanol or sucrose reinforcer.

## Introduction

Alcohol is one of the most widely used addictive substances. In the United States, the most recent estimates indicate that almost 50% of individuals aged 12 or older drank alcohol in the past month with approximately 22% of individuals in the same age range reporting at least one binge drinking episode during the same time frame (Substance Abuse and Mental Health Services Administration, 2023). In addition, recent data suggests that the prevalence of alcohol use disorder (AUD) is on the rise with an estimated 12.7% of individuals surveyed meeting criteria for diagnosis in 2012-2013 compared to 8.5% in 2001-2002 (Grant *et al*., 2017). Individuals with AUD experience a remarkably high burden of disease associated with a significant increase in disability and a high degree of mortality (GBD 2016 Alcohol Collaborators, 2018; Carvalho *et al*., 2019). Collectively, these data underscore the importance of research aimed at AUD prevention and treatment.

Rodent models are incredibly useful for studying the behavioral, pharmacological, and neurobiological underpinnings of AUD. Operant ethanol self-administration paradigms are particularly valuable as a method to interrogate the mechanisms underlying drinking behavior and assess the efficacy of pharmacotherapeutics to reduce ethanol intake. Despite its broad use in the field, most rodents do not readily self-administer large quantities of ethanol. Consequently, a considerable amount of effort has been put forth to identify methods that facilitate the highest possible levels of ethanol intake. Perhaps the most well-known effort in this direction is the sucrose-fade procedure pioneered by Samson et al. (1988). Using this approach, rats are first trained to self-administer a sucrose solution followed by introduction of ethanol to the solution. Over time the concentration of sucrose is progressively reduced until the reinforcing solution consists entirely of unadulterated ethanol. Although well-accepted by the field following its development, researchers eventually began to favor alternative approaches that avoided the potential confound introduced by the effects of a highly palatable reinforcer on mechanisms of reward and reinforcement that overlapped with those associated with ethanol. One alternative that has garnered relatively strong acceptance has been to habituate rats to ethanol prior to operant training using an intermittent-access two-bottle choice home cage drinking procedure. Using this approach, researchers showed that Long-Evans (Simms *et al*., 2010) and Sprague Dawley (Bito-Onon *et al*., 2011) rats will operantly self-administer ethanol without requiring the addition of sucrose to the reinforcer solution. Surprisingly, however, these studies did not include comparisons to a group that was naïve to ethanol prior to operant training making it impossible to determine the advantage of this approach over one that trains rats to operantly self-administer in the absence of prior experience drinking ethanol. Importantly, recent data suggests that the level of ethanol intake during intermittent-access two-bottle choice home cage drinking is not predictive of intake during operant self-administration (Patwell *et al*., 2021) adding further uncertainty to the benefit of this experimental approach.

More recently, several groups have suggested that shorter operant session duration can induce greater levels of ethanol consumption and, in particular, a high degree of binge-like drinking (Jeanblanc *et al*., 2018; Flores-Bonilla *et al*., 2021). Although exciting, it should be noted that both of these studies suggested that differences in drinking patterns contributed to changes in ethanol intake during operant sessions of shorter duration, yet, neither study used methods that incorporate direct measures of consumption. Use of such methods in studies of ethanol drinking patterns is crucial in light of work showing that a significant number of animals refrain from drinking ethanol despite completion of the operant response requirement (Blegen *et al*., 2018; Patwell *et al*., 2021). Thus, additional studies are needed, particularly those that measure consumption directly, in order to lend further support to this approach.

The present study was designed to directly assess the impact of prior experience with ethanol and changes in session duration on the acquisition and maintenance of operant ethanol self-administration in adult male and female Long-Evans rats. Our findings suggest that prior home cage drinking and shorter session duration are ineffective at facilitating higher acquisition rates or greater binge-like drinking patterns during operant sessions.

## Methods

### Animals

Adult male and female Long Evans rats (Envigo, Indianapolis, IN) were P60 upon arrival and allowed to acclimate to the vivarium for at least one week following shipment. Rats were singly housed in standard plexiglass cages under a 12 h reverse light-dark cycle (lights on at 22:00) and provided Teklad 7912 standard chow (Envigo) and water *ad libitum* throughout the duration of the study.

### Intermittent-access two-bottle choice home cage drinking

To determine the effect of prior experience drinking ethanol on the acquisition of operant ethanol self-administration, male rats were habituated to ethanol drinking in the home cage for three weeks prior to operant training. Rats assigned to the Water/Ethanol (W/E) group (n=24) underwent a standard intermittent-access two-bottle choice procedure identical to previously published procedures (Simms *et al*., 2008; Patwell *et al*., 2021). Rats were provided with one bottle of water and one bottle containing ethanol (20% v/v) in tap water on Mondays, Wednesdays, and Fridays approximately 30 min after the onset of the dark cycle. Change in bottle weight was recorded 30 min and 24 hours after session start. On all other days, rats were provided with two water bottles. Rats assigned to the Water/Water (W/W) group (n=24) were treated identically except that they received two bottles containing water seven days a week. Bottle changes on an empty cage were used to account for spillage.

### Operant self-administration

Operant training was performed in lickometer-equipped operant boxes (Med Associates, St Albans, VT) using previously published methods (Patwell *et al*., 2021). Rats were trained to self-administer either ethanol (20% v/v) or sucrose (1% w/v). To acclimate to the operant setting, rats first underwent 1-3 3-hour sessions where the sipper tube containing the reinforcer solution was permanently extended into the operant box. This was followed by 1-3 60-min magazine training sessions during which the sipper was non-contingently extended into the operant box for 30 sec at 30-sec intervals. Rats were then trained on a fixed ratio 1 (FR1) schedule of reinforcement during which a single lever press on the active lever resulted in illumination of a cue light above the active lever and extension of the sipper tube into the operant box for 15 sec. Sessions were 60 min in length and the location of the active lever was counterbalanced across rats. After ∼15 sessions, session duration was decreased to 30 min. After an additional ∼12 sessions the response requirement was increased to FR3. The ability of shorter session duration to increase intake was examined by reducing session duration to 15 min after rats had maintained FR3 responding during 30 min sessions for at least 10 days. Lever presses and licks on the sipper tube were recorded using MED-PC V software (Med Associates). Reinforcer-containing bottles were weighed before and after each operant session to calculate total fluid consumed. Rats were weighed daily following completion of each operant session in order to calculate intake in terms of body weight. All operant sessions began during the first three hours of the light-dark cycle.

### Blood ethanol concentration

Blood samples were collected via tail nick from a subset of male and female rats that successfully acquired operant ethanol self-administration for the purposes of analyzing blood ethanol concentration (BEC). Approximately 40 uL of blood was collected from the tail using a heparinized capillary tube immediately upon completion of the operant session. After collection, blood samples were centrifuged (10,000 RPM, 10 min, 4°C). The resulting plasma was collected and stored at - 20°C until ready for analysis using an AM1 Alcohol Analyzer (Analox Instruments, Ltd, Stourbridge, United Kingdom).

### Statistical analysis

Successful acquisition of operant ethanol self-administration was defined according to our previously published metrics with a “drinker” (Patwell *et al*., 2021) defined as a rat completing sufficient operant responses to achieve at least two reinforcer deliveries (i.e., sipper access) and consumption of greater than 0.1 g/kg ethanol on average per operant session. For operant self-administration, responding during the last five sessions at each duration were averaged together and used to compare behavior during 30-min and 15-min sessions. Drinking patterns were assessed by comparing operant data binned into 5 min increments across the session. Where applicable, paired two-tailed Student’s t-test and two-way repeated measures (RM) and mixed effects ANOVAs were used for statistical analyses. Males and females were run consecutively, not concurrently, preventing direct statistical comparisons between the sexes. All analyses were performed in GraphPad Prism Version 9 (San Diego, CA, USA).

## Results

### No effect of prior ethanol drinking on acquisition or magnitude of operant ethanol self-administration

Previous research suggested that prior experience with ethanol via intermittent access home cage drinking was sufficient to facilitate acquisition of operant ethanol self-administration in rats (Simms *et al*., 2010; Bito-Onon *et al*., 2011). However, whether this procedure increases acquisition rates over the absence of any initiation procedure has not been examined directly. To explore this, we compared operant performance in male rats that had intermittent access to ethanol in the home cage with those that had access only to regular drinking water prior to operant training (**Figure 1A**). Comparison of total fluid intake during home cage drinking using a mixed-effects two-way ANOVA revealed a main effect of drinking days [F(2.519,115.2)=8.787; p<0.0001] and a significant day by group interaction [F(8,366)=4.008; p=0.0001]. However, post-hoc comparisons failed to uncover any statistically significant differences in total fluid intake on any given day between rats in the Water/Water (W/W) group and rats in the Water/EtOH (W/E) group (all p values > 0.2; Sidak correction; **Figure 1B**). Similar to previous reports (Simms *et al*., 2010; Nentwig *et al*., 2019; Patwell *et al*., 2021), rats in the W/E group increased their ethanol intake progressively over the three-week home cage drinking period such that by the third week of drinking average 24 hr intake was 2.37 ± 0.40 g/kg (range: 0.00-5.73 g/kg; **Figure 1C**).

**Figure 1.**
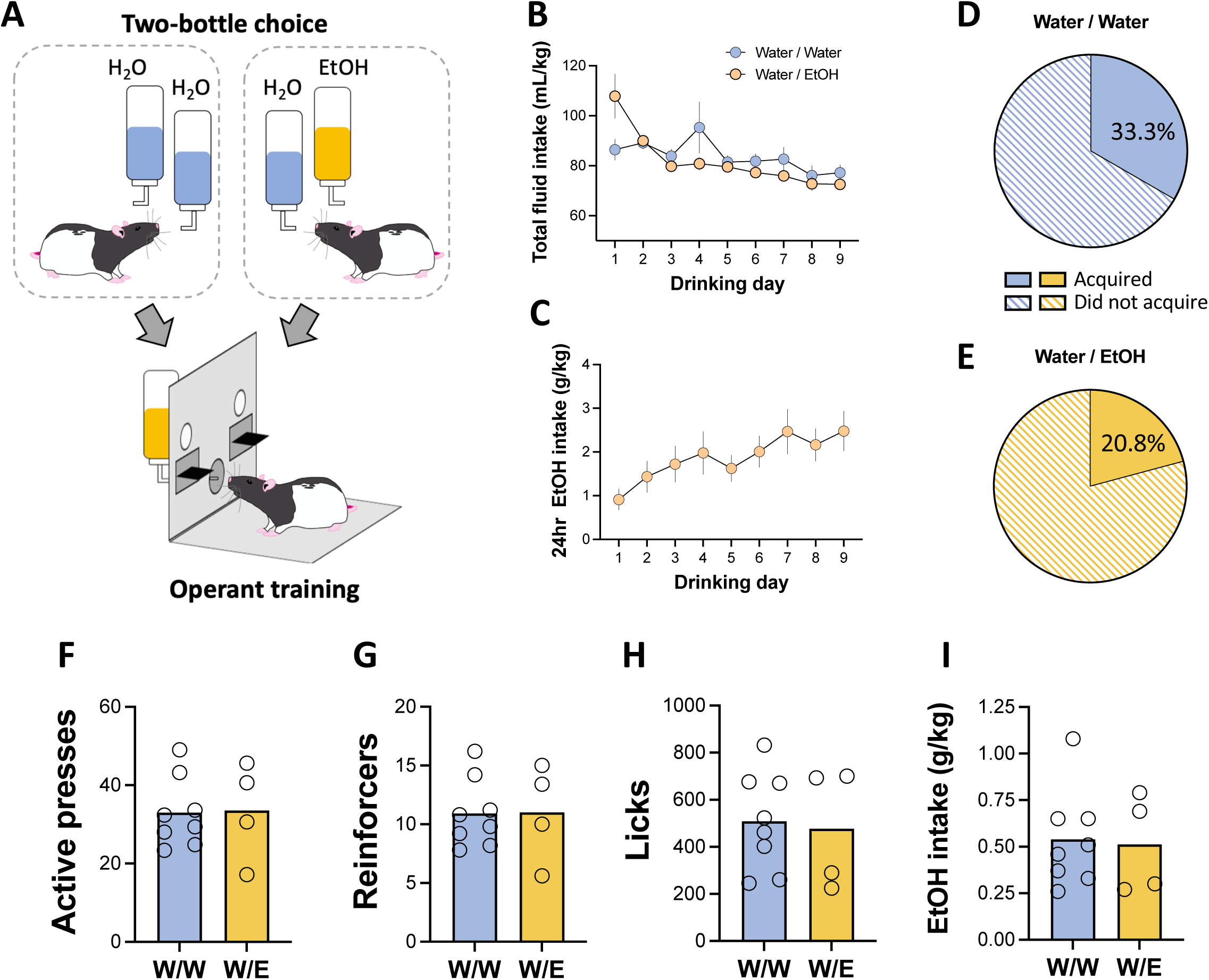
Home cage access to ethanol does not enhance acquisition or magnitude of operant ethanol self-administration in male Long-Evans rats. **(A)** Comparisons were made between rats that received intermittent-access to one bottle of water and one bottle of ethanol or two bottles of water prior to operant training (n=24/group). **(B)** Total fluid intake was not significantly different between W/W and W/E groups. **(C)** Ethanol intake increased over drinking days in agreement with previous reports. Acquisition rates did not differ between W/W **(D)** and W/E **(E)** groups. In addition, neither appetitive responding **(F-G)** or the magnitude of intake **(H-I)** differed as a result of prior history with ethanol.

After ∼5 weeks of operant training, approximately 27% of all rats had successfully acquired operant ethanol self-administration. A Fisher’s exact test comparing acquisition rates between groups showed that a similar proportion of rats in the W/W and W/E groups acquired operant self-administration (p=0.517; **Figure 1D-E**). Unpaired Student’s t-tests were used to compare operant performance between rats from each group that acquired self-administration. This analysis showed that rates of active lever presses (t=0.08451, df=10; p=0.934) and reinforcer deliveries (t=0.03660, df=10; p=0.972) were not significantly different between W/W and W/E groups (**Figure 1F-G**). Number of licks (t=0.2325, df=10; p=0.821) and total ethanol intake (t=0.1635, df=10; p=0.873) also did not differ significantly between groups (**Figure 1H-I**). Together, these data suggest that home cage ethanol drinking prior to operant training has no effect on acquisition rates or overall magnitude of appetitive or consummatory responding during operant ethanol self-administration sessions.

### Ethanol intake is reduced when operant session duration is shortened

Recent work suggests that shortening operant session duration can facilitate higher rates of responding and greater binge-like ethanol intake (Jeanblanc *et al*., 2018; Flores-Bonilla *et al*., 2021). However, these studies inferred ethanol consumption from appetitive responding. To address this gap, we compared operant performance in male and female Long-Evans rats using a lickometer-equipped system that directly measures ethanol drinking. Male rats that underwent intermittent-access two-bottle choice home cage drinking prior to operant training were transitioned from 30-min sessions to sessions lasting 15-min. Data from these rats were combined with additional data from male and female rats that did not undergo home cage drinking (since prior experience with ethanol had no significant impact on acquisition of operant self-administration).

Both males and females successfully discriminated between active and inactive levers. This was confirmed using paired Student’s t-tests, which showed that rats made significantly more active than inactive lever presses during both 30-min (male: t=14.32, df=18, p<0.0001; female: t=4.693, df=6, p=0.0034) and 15-min (male: t=8.741, df=18, p<0.0001; female: t=6.593, df=6, p=0.0006) sessions (data not shown). Next, paired Students t-tests were used to compare operant performance within each sex across 30-min and 15-min sessions. In males, this analysis revealed a significant decrease in measures of both appetitive and consummatory behavior. As shown in **Figure 2A-D**, active presses (t=4.410, df=18; p=0.0003), reinforcers delivered (t=4.284, df=18; p=0.0004), licks (t=3.429, df=18; p=0.0030), and intake (t=3.309, df=18; p=0.0039) were all significantly lower during 15-min than 30-min sessions.

**Figure 2.**
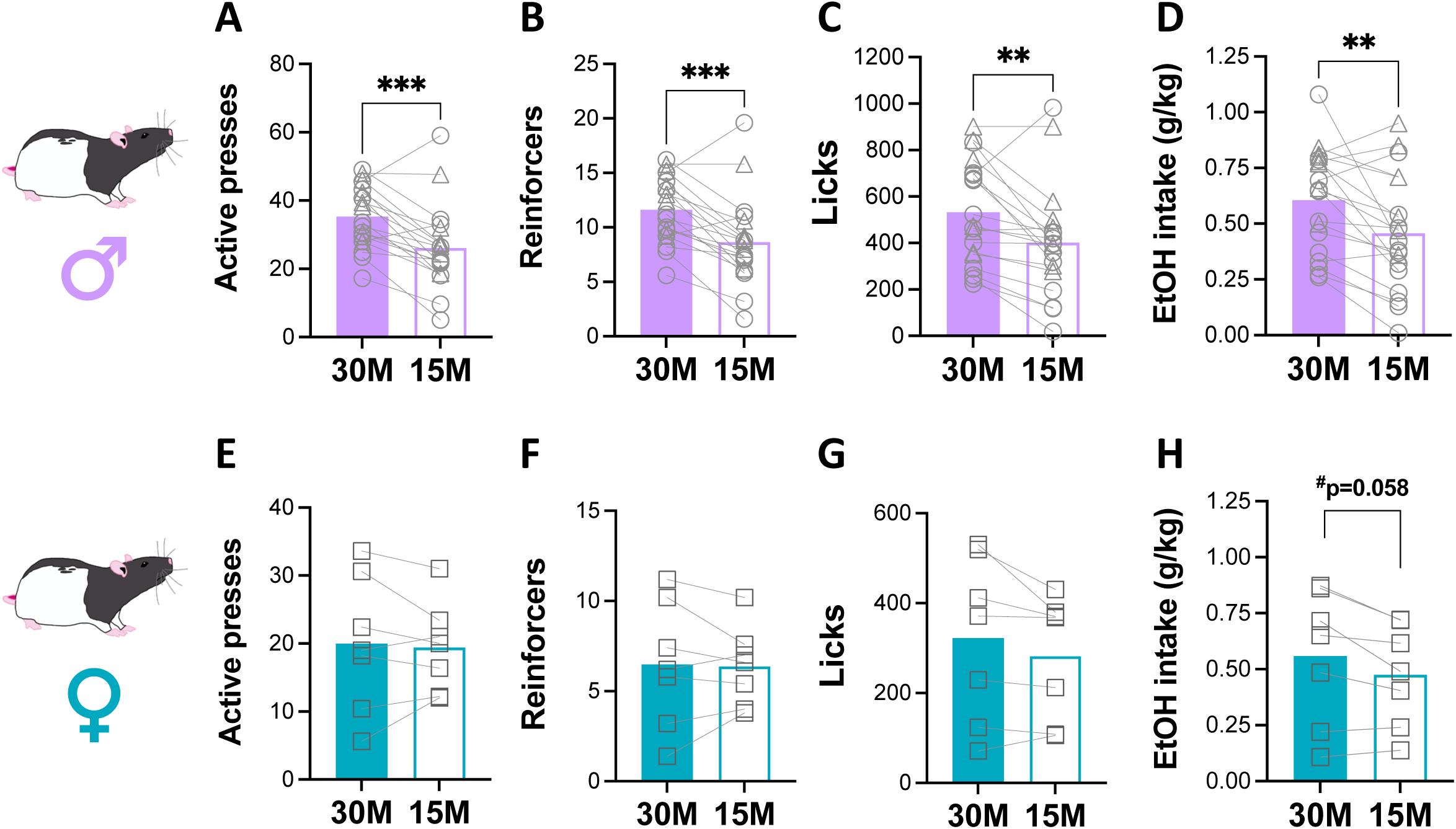
Shorter session duration did not increase appetitive or consummatory behaviors during operant ethanol self-administration. In males **(A-D)**, shortening the operant session duration resulted in a significant decrease in both appetitive and consummatory behaviors. This was evident in the number of active lever presses **(A)** and number of reinforcers acquired **(B)** as well as the number of licks **(C)** and total ethanol consumed **(D)**. In contrast, session length had no effect on the number of active lever presses **(E)** or reinforcers delivered **(F)** in females. While females did not exhibit a significant difference in the number of licks **(G)** between sessions of different durations, they did exhibit a slight reduction in total ethanol intake **(H)** in 15-min compared to 30-min sessions, which trended toward statistical significance. In A-D, circles correspond to rats that underwent two-bottle choice prior to operant training as in Figure 1; triangles correspond to rats that did not participate in this experiment and therefore had no experience with ethanol prior to operant training. **p≤0.01, ***p≤0.001.

Interestingly, shorter session duration was not associated with a similar decrease operant performance in females (**Figure 2E-H**). In fact, the number of active lever presses (t=0.3299, df=6; p=0.7527) and reinforcer deliveries (t=0.1881, df=6; p=0.8570) did not differ significantly between 30-min and 15-min sessions. Number of licks and ethanol intake were slightly, but not significantly, lower in 15-min compared to 30-min sessions, although this effect trended toward significance for intake (licks: t=1.738, df=6, p=0.1329; intake: t=2.330, df=6; p=0.0586). Altogether, these results suggest that shorter operant sessions do not enhance operant performance or increase total ethanol intake in male and female rats.

### Rate of ethanol intake is unaffected by shorter operant session duration

Previous work suggests that shorter session duration increases the rate of ethanol intake thereby producing greater levels of intoxication than is observed in sessions of longer duration (Jeanblanc *et al*., 2018). Therefore, we next considered the possibility that while shorter operant sessions did not increase overall levels of ethanol intake, they may have facilitated an increase in rate of consumption. To examine this, we assessed the pattern of ethanol intake by comparing the average number of licks occurring in 5-min bins across 30-min and 15-min sessions. While a two-way RM ANOVA of drinking patterns in males uncovered a trend for a significant main effect of bin [F(1.1,12)=4.3, p=0.0580], no significant main effect of session duration [F(1.0,11)=1.3, p=0.2796] or interaction between the two factors [F(1.4,16)=0.76, p=0.4416] was observed (**Figure 3A**). Interestingly, while the same analysis in females found no main effect of bin [F(1.167,7.000)=3.067, p=0.1213] or interaction between bin and session duration [F(1.515,9.092=1.653, p=0.2407], it did reveal a main effect of session duration [F(1,6)=8.331, p=0.0278] with significantly greater licks occurring during the first three 5-min bins of 15-min sessions compared to 30-min sessions (**Figure 3E**). It should be noted, however, that although statistically significant, this difference in average lick number was only modest (30-min: 81.93 ± 7.984; 15-min: 93.04 ± 18.16). Cumulative records from representative rats further revealed that drinking patterns were largely similar during the first 15 min of all operant sessions. For example, lick pattern and overall intake did not differ in a subset of male (**Figure 3B**) and female (**Figure 3F**) rats whose self-administration was restricted to the first 15 min of the operant session regardless of session duration. Other rats exhibited relatively similar patterns of intake during the first 15 min of the session but then had a large drinking bout (**Figure 3C, 3G**) or several small drinking bouts (**Figure 3D, 3H**) in the second half of 30-min sessions, which served to increase overall intake in sessions of longer duration. In addition, no significant difference in BEC (t=0.9209, df=9; p=0.3811) was observed in blood samples taken from a subset of male and female rats immediately after completion of 30-min and 15-min sessions (**Figure 4**). Altogether, these data suggest that shortening operant session duration does little to promote binge-like ethanol drinking.

**Figure 3.**
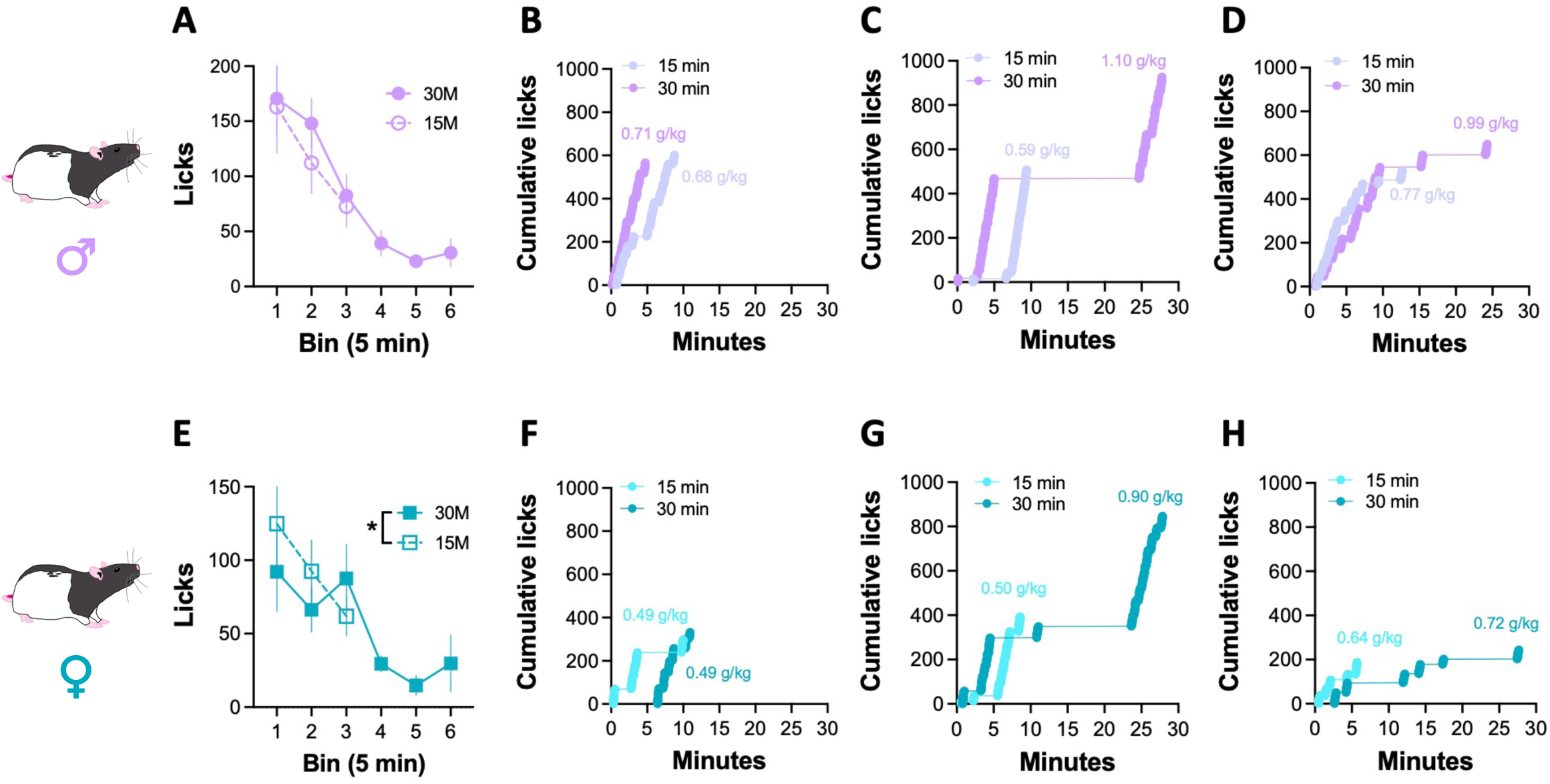
Little-to-no impact of shorter session duration on ethanol drinking patterns. Average lick pattern for ethanol was not significantly different in males **(A)** during first three 5-min bins of the operant session regardless of whether the session lasted 15 or 30 min in duration. This is evident in representative cumulative records, which show that intake during the first 15 min of the operant session was unchanged after session length was reduced. This resulted in similar levels of intake in rats whose intake was restricted to the first 15 min of the session regardless of session length **(B)** but reduced intake in rats that drank considerable **(C)** or relatively small **(D)** amounts of ethanol during the final 15 min of 30-min sessions. In contrast to males, females exhibited a slight, but significant, increase in average licks during the first three 5-min bins of the operant session **(E)** when session duration was shortened from 30 to 15 min. Although statistically significant, representative cumulative records suggest that this modest increase in licks was insufficient to increase intake. Indeed, similar to males, a subset of females exhibited similar levels of intake during 30- and 15-min sessions **(F)**, whereas others achieved lower levels of intake as a result of missed opportunity for additional large **(G)** and small **(H)** drinking bouts during longer duration sessions.

**Figure 4.**
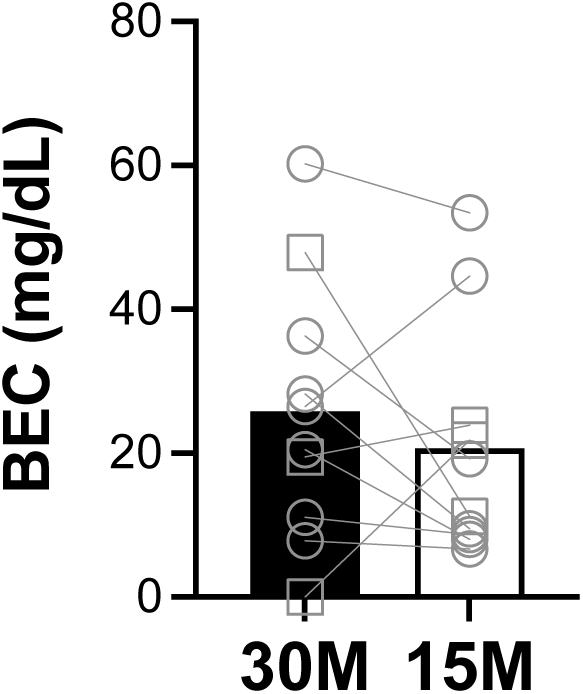
BECs are unaffected by session duration. Male (circles; n=7) and female (squares; n=3) rats achieved similar BECs during 30-min and 15-min sessions.

### Shortening session duration also decreases appetitive and consummatory responses for sucrose

To determine whether the effect of shorter session duration on operant self-administration was specific to ethanol, we next examined operant performance across 30-min and 15-min drinking sessions using sucrose as the reinforcer. Paired Student’s t-tests confirmed that both males and females successfully discriminated between the active and inactive lever throughout testing [males: 30 min (t=7.082, df=7, p=0.0002); 15 min (t=7.001, df=7, p=0.0002); females: 30 min (t=2.774, df=6, p=0.0323); 15 min (t=2.524, df=6, p=0.0450). Similar to what we observed in rats trained to self-administer ethanol, male rats trained to operantly self-administer sucrose exhibited a reduction in both appetitive and consummatory behavior when session duration was shortened from 30 to 15 min (**Figure 5A-D**). This was evident from results of paired Student’s t-tests, which showed significantly fewer active lever presses (t=3.648, df=7, p=0.0082), reinforcer deliveries (t=3.559, df=7, p=0.0092), licks (t=2.454, df=7, p=0.0438), and overall intake (t=2.515, df=7, p=0.0401). Similar to males, females trained to lever press for sucrose also exhibited a significant decrease in operant performance for sucrose when session duration was decreased from 30 to 15 min (**Figure 5F-I**). Indeed, paired Student’s t-tests found significantly fewer active lever presses (t=3.217, df=6, p=0.0182), which corresponded to fewer reinforcer deliveries (t=2.894, df=6, p=0.0275). This was paired with significantly fewer licks (t=2.626, df=6, p=0.0393) and reduced overall intake (t=2.780, df=6, p=0.0320).

**Figure 5.**
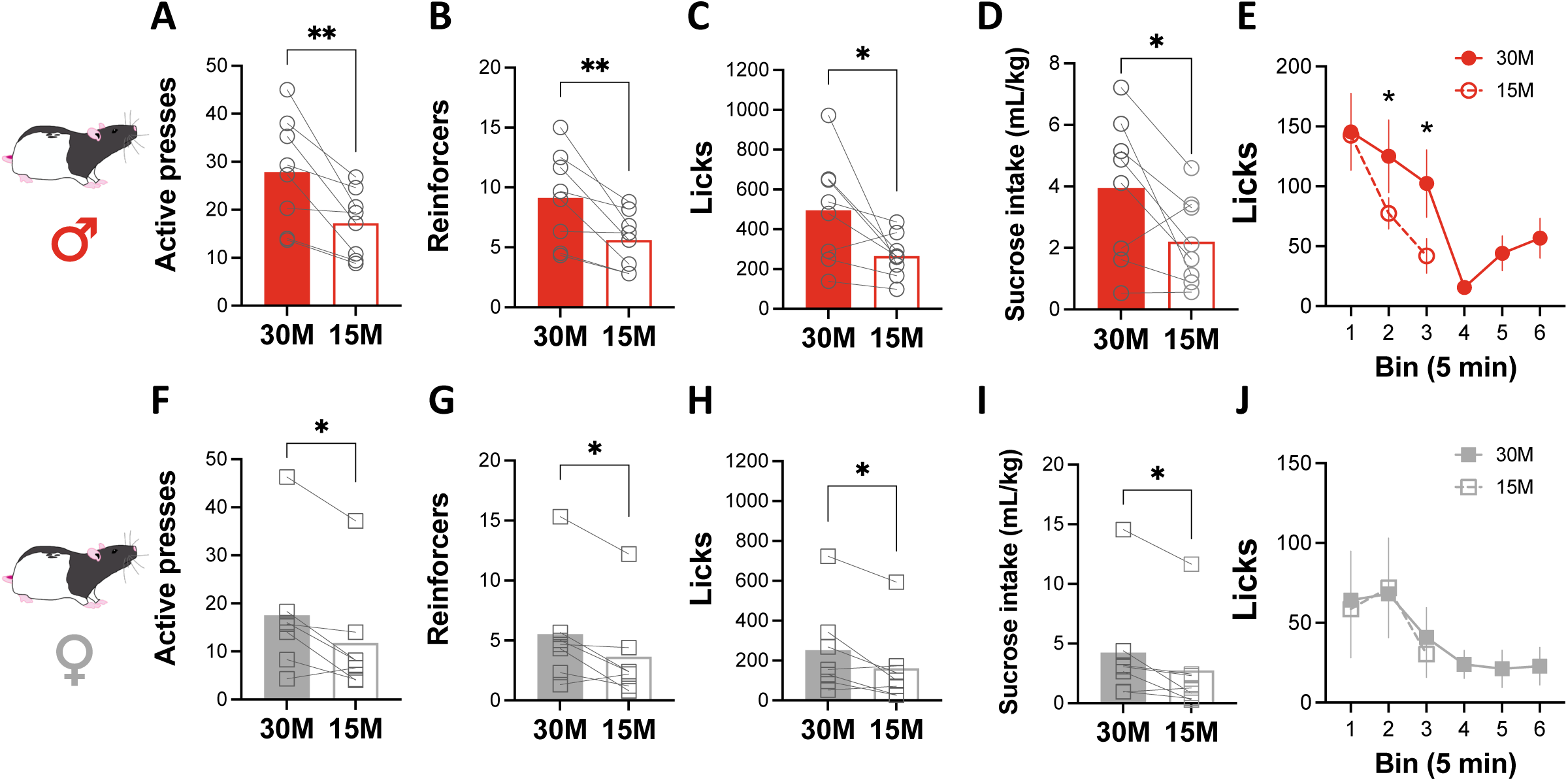
Shortening session duration decreased appetitive & consummatory behaviors during operant sucrose self-administration. In males, shortening operant session duration resulted in a significant decrease in the number of active lever presses **(A)** and reinforcers earned **(B)** as well as the number of licks **(C)** and overall sucrose intake **(D)**. The decrease in intake in males was driven by significantly fewer licks during the last two-thirds of the 15-min operant session **(E)**. Shortening session duration produced similar results in females who performed significantly fewer active lever presses **(F)** corresponding to fewer reinforcer deliveries **(G)** as well as a slight, but significant decrease in licks **(H)** and overall intake **(I)** in 15-min compared to 30-min sessions. In contrast to males, however, changes in total licks were sufficiently modest that they had no effect on overall drinking pattern **(J)**. *p≤0.05; **p≤0.01.

We next assessed within-session dynamics to determine whether the observed change in sucrose self-administration was driven by a change in drinking pattern. In males, a two-way RM ANOVA examining average licks during 5-min bins across the operant session found a significant main effect of bin [F(2,14)=6.211, p=0.0117] but no main effect of session duration [F(1,7)=2.072, p=0.1932]. This analysis also uncovered a significant interaction between bin and session duration [F(2,14)=4.138, p=0.0387]. Posthoc comparisons using a Sidak correction found that rats performed significantly fewer licks in the second (p=0.0188) and third (p=0.0035) 5-min bin during 15-min sessions compared to 30-min sessions (**Figure 5E**). These results suggest that shorter session duration promotes reduced engagement shortly after session start in males. Unlike in males, the same analysis in females found no significant main effects of either bin [F(2,12)=3.037, p=0.0857] or session duration [F(1,6)=0.1238, p=0.7370] or an interaction between the two factors [F(2,12)=0.2832, p=0.7583]. As shown in **Figure 5J**, these data suggest that for females, lost drinking opportunity during the second half of the session plays a greater role in driving the reduction in sucrose intake occurring during shorter operant sessions.

## Discussion

Despite studies suggesting that exposure to ethanol using an intermittent-access two-bottle choice procedure can facilitate acquisition of operant ethanol self-administration, no study has directly compared acquisition rates between rats that are ethanol-exposed and ethanol-naïve prior to operant training. Unexpectedly, results from the present study found that acquisition rates were similar between adult male Long-Evans rats that did and did not have prior experience with ethanol drinking. Moreover, operant performance was similar between groups once rats were fully trained further indicating that prior ethanol exposure does not facilitate greater levels of intake in the operant paradigm. These data are in general agreement with our previous work, which showed that the amount of ethanol consumed in the home cage was not predictive of consumption in the operant setting (Patwell *et al*., 2021).

Using the same definition of successful acquisition as was used in the present study, our previous work found that approximately 50% of male Long-Evans rats acquire operant self-administration (Patwell *et al*., 2021). In contrast, acquisition rates were substantially lower (∼27%) in the current study. Construction in the building housing the animals may have contributed to these lower rates. However, rats from both groups were run concurrently and therefore would have been equally influenced by the construction. Importantly, our assessment of the impact of prior home cage ethanol drinking on operant ethanol self-administration is limited to male rats. While we cannot make conclusions about the impact of this approach on operant performance in female rats, our ongoing work suggests that a similar disconnect exists between home cage drinking and operant self-administration in females (unpublished observations).

Unlike previous work, results from the present study do not support the notion that shorter operant sessions facilitate greater ethanol intake and promote binge-like drinking patterns. Instead, we found that consumption is significantly lower during shorter sessions in males. Inspection of drinking patterns enabled with the use of a lickometer-equipped system revealed that this was due, in large part, to missed drinking bouts that the rats typically performed in the second half of the 30-min operant session. Interestingly, female rats did not exhibit the same decrease in behavior observed in males during shorter operant sessions. Instead, appetitive behaviors were similar between long and short duration sessions. Females also exhibited a similar degree of consummatory behavior independent of differences in session duration, although a trend toward a decrease in total ethanol intake was observed in 15-min compared to 30-min sessions. Although non-significant, these data further suggest that reducing the operant session from 30-min to 15-min is not necessarily an advantageous approach for researchers interested in modeling high levels of ethanol consumption.

Methodological differences could contribute to the discrepant findings between the present study and the two previous studies examining the effect of session duration on ethanol self-administration. For example, Jeanblanc et al. (2018) assessed drinking using a between-subjects design that compared intake in “heavy” drinking rats trained to self-administer in 30-min sessions to “binge-like” rats that transitioned to 15-min sessions after being trained in a manner identical to “heavy” drinking rats. While this study makes a compelling case for shorter session duration facilitating increased drinking rates and higher total ethanol consumption, it remains possible that cohort differences drove the observed effects. Within-subject comparison of operant performance during 30- and 15-min sessions within the “binge-like” group could help to address this concern. It should also be noted that operant testing in the Jeanblanc et al. (2018) study was performed during the light portion of the light-dark cycle, whereas rats in the current study were tested during the dark cycle – a time point that corresponds to normal waking hours for rodents. This methodological difference could also contribute to conflicting findings between the two studies, particularly in light of findings from Flores-Bonilla et al. (2021), which showed that changes in session duration affected operant performance only at certain times of the light-dark cycle. Still, our findings disagree with those of Flores-Bonilla et al. (2021) despite testing conditions used in the present study closely paralleling theirs (e.g., testing during beginning of dark cycle). Notably, unlike in the present study, Flores-Bonilla et al. (2021) inferred consumption from appetitive responding. Although the authors went to considerable lengths to confirm that appetitive responding accurately reflected consumption by inspecting the operant apparatus for spillage and unconsumed ethanol, it remains possible that findings would be different with the use of more direct measures of drinking.

Of significance, we showed a similar effect of session duration on consumption of sucrose reinforcer in separate cohorts of male and female rats. At first glance, reduced sucrose intake during shorter sessions may not be particularly surprising since rats are not consuming sucrose for its pharmacological effects. However, drinking patterns in these rats closely parallel those observed when ethanol is used as the reinforcer with the majority of intake occurring during the first half of a 30-min session. Thus, these data lend further support to our conclusion that shorter session duration does not always promote increased reinforcer consumption. Altogether, our findings and those from previous studies (Jeanblanc *et al*., 2018; Flores-Bonilla *et al*., 2021) point to the need for more work examining the role that session duration may or may not play in promoting high levels of ethanol intake in the operant setting.

While historically the field has relied primarily on drinking data derived from males, an increasing number of studies report data from animals of both sexes. Much of this work has shown that female rats consume greater quantities of ethanol than males in home cage drinking paradigms (see Priddy *et al*., 2017 for review). However, far fewer studies have examined sex differences in the operant setting. The few that have, report mixed results with females drinking more (Blanchard *et al*., 1993) or equal (Moore and Lynch, 2015; Priddy *et al*., 2017) amounts to males. Male and female rats in the present study were run consecutively, not concurrently, preventing us from being able to make direct statistical comparisons of sex differences in operant performance. Nevertheless, visual inspection of the figures included in the present study suggest that male and female rats consume similar amounts of ethanol during 30-min sessions but that females perform fewer appetitive responses. This is further supported by descriptive statistics revealing a similar degree of ethanol intake (males: 0.61±0.05; females: 0.56±0.11) but fewer appetitive responses (males: 35.27±2.10; females: 19.97±3.80) and fewer licks (males: 532.4±50.51; females: 322.3±69.61) in females than in males. While inclination may be to suggest that these differences reflect a more efficient drinking strategy by females, it should be noted that females require less volume (mL) to achieve the same level of ethanol intake (g/kg) due to their lower body weight (kg) compared to males. Therefore, it is not unexpected that females would perform fewer licks (i.e., obtain less volume) in order to achieve the same level of intake as males. Indeed, comparison of average licks performed per reinforcer (i.e., sipper) access (males: 45.07±2.74; females: 43.57±4.09) shows that male and female rats in the present study are likely using similar behavioral strategies to consume similar quantities of ethanol in the operant setting. Similar patterns of intake were observed in male and female rats self-administering sucrose, with females gaining access to the reinforcer fewer times and performing fewer licks but performing a similar number of average licks per reinforcer access (males: 52.57±4.81; females: 43.57±4.09) to achieve relatively similar levels of sucrose intake to males (males: 3.94±0.83; females: 4.25±1.78). Altogether, these data reveal a similar degree of operant ethanol and sucrose self-administration in adult Long-Evans rats of both sexes.

In summary, the present study found that intermittent-access two-bottle choice ethanol drinking provided no advantage over home cage access to regular drinking water as an initiation procedure that has previously been suggested to enhance acquisition of operant ethanol self-administration. We further showed that shortening operant session duration was ineffective at increasing overall ethanol intake or in promoting greater binge-like drinking patterns. These findings should be taken into consideration as researchers design experiments using operant ethanol self-administration to examine ethanol drinking behavior in adult male and female rats.

## Data availability statement

Data are available upon request.

